# Concurrent presentation of memory-related odors and sounds nullified sleep reactivation benefits

**DOI:** 10.1101/2025.07.25.666857

**Authors:** Gayathri Subramanian, Christina Zelano, Ken A. Paller, Eitan Schechtman

## Abstract

Reactivation of recently acquired memories during sleep supports their longevity. Reactivation can be altered during sleep using odors or sounds through a technique termed targeted memory reactivation (TMR). Here, we attempted to selectively weaken memories by reactivating them together with forgetting instructions. We delivered sounds to reactivate spatial memories and concurrent odors to reactivate instructions. Participants learned about the instructions in a Directed-Forgetting task performed with a list of to-be-remembered and to-be-forgotten words. One odor was linked with instructions to forget, one with instructions to remember, and a third was not assigned any meaning. During a nap, sounds previously linked with object-location learning were simultaneously presented with these odors. Spatial recall was tested after sleep. Sound cues produced a selective recall benefit for weakly encoded memories. However, memory results did not support the prediction that forgetting could be instilled by the concurrent forget odor. An encoding-strength-dependent benefit was largest when sounds were presented together with the odor that lacked assigned meaning, whereas the other two odors both disrupted sound-induced memory reactivation. These two odors were linked with instructions and with multiple learning episodes in the Directed-Forgetting task. Accordingly, we infer that reactivation evoked by the remember and forget odor cues interfered with the reactivation of spatial memories. Odors also induced a prolonged decline in sigma EEG power (12–16 Hz) that continued at least 10 s after odor offset. Overall, these findings highlight the complexity of memory consolidation during sleep when multiple memories and multiple cues are involved.

## Introduction

Sleep potentially influences memory in multiple ways (Poe, 2017; Stickgold, 2005). A pivotal question with both theoretical and practical implications is whether sleep manipulations can systematically weaken memories during sleep. Such manipulations, if operative reliably, could be of practical usefulness. For example, noninvasively weakening memories during sleep might help some patients suffering from disorders such as post-traumatic stress disorder. Furthermore, memory consolidation is thought to fundamentally involve interrelationships among memories where some memories or memory features might be strengthened while others are weakened.

A prominent way to experimentally alter memory storage during sleep is known as Targeted Memory Reactivation (TMR). TMR involves presenting learning-related stimuli unobtrusively during sleep to reactivate memories (Hu et al., 2020; Oudiette & Paller, 2013). TMR has been shown to boost memory retention using sounds or odors. However, in a few studies, TMR has also been employed to induce forgetting and weaken memories. For instance, consolidation during sleep was shown to be disrupted by playing learning-related sound cues at a high volume to cause arousals, reducing memory recall after sleep (Whitmore et al., 2024; Whitmore & Paller, 2023). Employing another approach, memories were shown to be weakened through reactivation-induced forgetting when a cue common to overlapping memories was presented during sleep (Joensen et al., 2022).

For a memory-weakening intervention to be clinically viable, the risk of accidentally strengthening a traumatic memory should be extremely small. To mitigate this risk, other approaches have avoided presenting cues directly linked with the to-be-weakened memory, and turned instead to approaches building on wake-based methods for motivational forgetting. One such approach for weakening memories during sleep made use of a Directed-Forgetting task, whereby participants were instructed to either commit information to memory or try to avoid doing so (Woodward & Bjork, 1971). In a TMR study building on Directed-Forgetting literature, Schechtman and colleagues (2020) required participants to learn object locations associated with sounds that conveyed instructions to either remember or forget object-related information. Playing a Forget sound during sleep enhanced forgetting for corresponding memories. This approach required co-activation of two memories during sleep: the to-be-forgotten memory and the instruction to weaken it. Reactivating the former independently would lead to strengthening; reactivating the latter independently would not impact the target memory. Schechtman and colleagues (2020) were able to reactivate both together by linking the same sound to both the forgetting instructions and the to-be-forgotten object location. However, this approach is not translatable: traumatic memories are rarely, if ever, associated with weakening cues during encoding.

Using another approach not subject to this issue, Simon and colleagues (2018) trained participants to associate a tone with forgetting using a Directed-Forgetting task. Then, participants completed a separate task, learning a set of 28 objects, each associated with a related sound. During sleep, object sounds for 5 objects were repeatedly played together with the forget tone, presented 3 s after the offset of the object sound each time. Seven days later, spatial free recall was impaired for the cued objects compared to control objects, suggesting that participants were able to conjointly process two unrelated stimuli during sleep to suppress subsequent memory. However, an alternative explanation is that the second sound did not induce forgetting as intended but rather had the generic effect of disrupting the consolidation process initiated by the object-related sound.

Indeed, other results suggest that consecutive presentations of sounds with a short inter-stimulus interval (ISI) during sleep blocks reactivation. In a study by Schreiner and colleagues (2015), participants learned Dutch-German word pairs. During sleep, Dutch words were presented, followed by their German counterpart (ISI = 200 ms). This cueing regimen did not produce any sleep-reactivation benefit. A similar study used ISIs of 1000–1500 ms and found a reactivation benefit only for ISIs between 1373–1500 ms (Farthouat et al., 2017). In both studies, shorter ISIs reduced EEG power in the theta (5–8 Hz) and sigma (11–16 Hz) bands, both of which are linked with memory processing during sleep. These studies highlight that the presentation of two auditory stimuli in close temporal proximity may interfere with reactivation and prevent behavioral effects (see also Lazarus et al., 2025). If interference was the cause of the forgetting effect observed in Simon et al. (2018), such a manipulation would not be effective in weakening well-established, salient memories, such as traumatic memories.

Here, to attempt to avoid interference between stimuli, we used a multimodal strategy to modify memories. We speculated that multimodal simultaneous presentation would more flexibly link the weakening cues to previously established memories, possibly enhancing the weakening effect and establishing the potential of sleep-dependent suppression. During a nap, odors and sounds were presented simultaneously. We speculated that forgetting could be triggered by an odor to weaken previously established memories linked with sounds.

## Materials and methods

### Participants

We recruited 52 participants from the community. Twenty-three participants were excluded because they did not reach stable non-rapid-eye-movement (NREM) sleep (N2 and N3 or slow-wave sleep, SWS) for cueing, two participants were excluded because of issues with their performance (see below), two participants were excluded due to technical failure of the EEG system, one participant withdrew because they had trouble hearing auditory stimuli, and one participant withdrew after experiencing discomfort due to odors. The exclusion rate was higher than that in most TMR studies because of longer cueing periods during sleep (with both odors and sounds, requiring 6–7 min), increasing the probability of arousals and insufficient cueing. The final sample of 23 participants (16 identified as women, seven identified as men, age = 22.52 ± 0.59 years, mean ± SE) had no known history of neurological, sleep, respiratory disorders, significant food or non-food allergy, smell, taste, or ear-nose-throat disorder, and sinusitis or allergic rhinitis. In preparation for the study, participants were asked to go to bed later than usual and wake up earlier than usual to limit their hours of sleep the night before. They were also asked to avoid consumption of caffeinated beverages or alcohol on the day of the study. The Northwestern University Institutional Review Board (IRB) approved the study.

### Materials

Visual stimuli were presented on a laptop (1920 × 1080 pixels) with sounds presented using external PC speakers. Visual stimuli consisted of 108 words randomly selected from a set of 175 words (2.01 ± 0.83 syllable length) taken from Wylie et al. (2008) and 60 images (120 x 120 pixels) taken from the Bank of Standardized Stimuli database (Brodeur et al., 2010). Images were accompanied by congruent sounds (e.g., an image of a cat along with *meow;* 0.52 ± 0.01 s, range of 0.17 s to 0.6 s). Similar sets of stimuli were used in other recent TMR studies (e.g., Lazarus et al., 2025; Schechtman et al., 2023). Odors were delivered via a rubber mask through a 12-channel, computer-controlled, air-dilution olfactometer with two mass flow controllers to allow for precisely timed and concentration-controlled stimuli. Odor bottles containing fresh banana, peanut butter, parmesan cheese, coffee beans, or soap were connected to the olfactometer and used as olfactory stimuli. The EEG signal was recorded from 32 electrodes using a BrainVision system on BrainVision Pycorder software. The stimuli and odors were controlled using Psychtoolbox-3 on MATLAB R2020b (MathWorks Inc., Massachusetts, USA).

### Procedure

The experimental procedure is depicted in Fig 1a. After consenting to participate, participants were fitted with the odor mask and were asked to smell five odors and pick the three they found most distinguishable. Participants were then told which odor was going to be the Remember odor and which the Forget odor. Participants then underwent a Directed-Forgetting Task, which was intended to solidify the odor associations. This task was followed by a Spatial Memory Task, wherein on-screen positions of 60 objects were learned. After a pre-sleep memory test on the spatial locations, participants were fitted with the EEG cap and were given a 90-min nap opportunity. During stable NREM sleep, odors and sounds were played simultaneously as described below. Upon waking up, participants completed post-sleep tests on both the Spatial Memory Task and the Directed-Forgetting Task.

**Figure 1.**
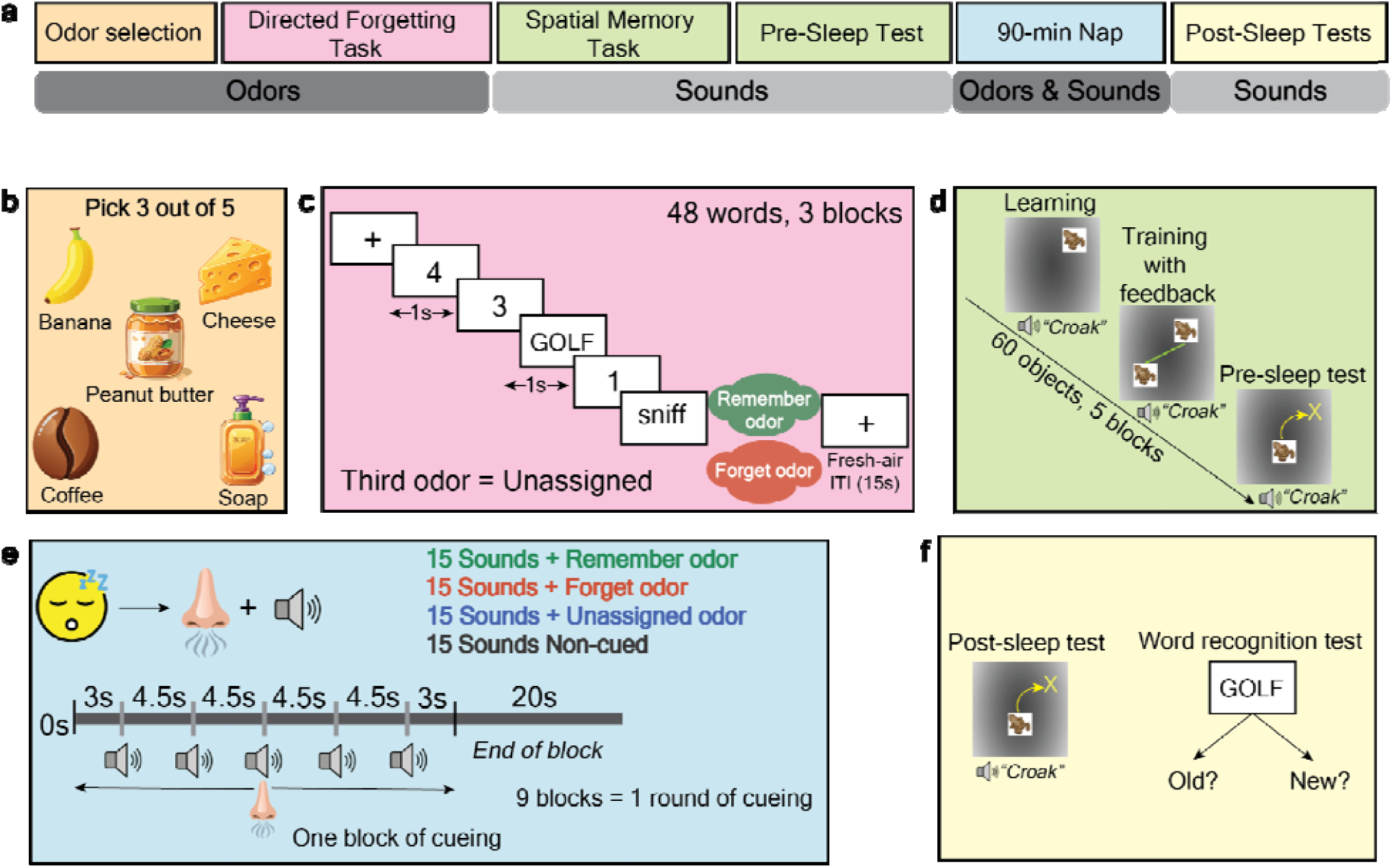
Experimental procedure **a** Overview of the experimental design. **b** Participants were presented with five odors and instructed to pick three that were most distinguishable. **c** In the Directed-Forgetting Task, two of the three odors were randomly assigned to the Remember and Forget conditions. Participants were instructed to remember or forget words based on which odor cue followed. They were exposed to 48 words split between three blocks. **d** The Spatial Memory Task involved a learning phase, a training phase with feedback, and a Pre-Sleep Test. This task included 60 objects learned over five blocks. **e** Participants were given the opportunity to nap for 90 min; when they entered stable NREM sleep, odors and sounds were presented. Each of the three odor conditions was paired with 15 sounds. The remaining 15 sounds were not presented during sleep (Non-cued). In each cueing block, the odor was presented throughout. 3 s after odor onset, five sounds were played consecutively (with 4.5 s onset-to-onset interval). **f** Post-Sleep tests involved both the Spatial Memory Task and a recognition test on the words from the Directed Forgetting Task.

### Odor selection

Participants were presented with five odors: banana, peanut butter, parmesan cheese, coffee beans, and soap (Fig 1b). Participants used a computer interface to control the odor presentation and were allowed to smell each one multiple times. As they smelled the odors, they were asked to name the odors and feedback was provided. They then chose three odors that they found to be distinct and recognizable. The banana, peanut butter, parmesan, coffee, and soap odors were chosen by 56.52%, 69.56%, 8.70%, 91.30%, and 73.91% of the participants, respectively. The remaining two odors were not used for the rest of the experiment.

After selecting the three odors, they were presented for 4 s each, and participants were asked to rate their intensity and pleasantness on a 7-point scale ranging from “not intense” to “very intense" and from “very unpleasant” to “very pleasant” respectively.

The three selected odors were randomly assigned to be the Remember condition, the Forget condition, and the Unassigned condition. The odors in the Remember, Forget, and Unassigned conditions had comparable pleasantness ratings (5.09 ± 0.23, 4.87 ± 0.28, and 5.30 ± 0.35, respectively) and intensity ratings (4.83 ± 0.29, 4.87 ± 0.27, and 4.70 ± 0.31, respectively).

### Odor Matching Task and Directed Forgetting Task

Two odors were linked with instructions to either remember or forget in a Directed Forgetting Task. For our study, we adopted the most commonly used version of this task, which involves instructions to remember or forget visually presented words (e.g., Santangelo et al., 2025; Woodward & Bjork, 1971). The instructions for this task were explained to the participants as follows. Words would appear on the screen followed by one of the two assigned odors (Fig 1c). They should try to remember the preceding word if they smell the Remember odor, and they should try to avoid registering the word if they smell the odor linked with forgetting. Before starting the Directed Forgetting Task, participants learned to match the odors to their respective instructions (i.e., Odor matching Task). They used a computer interface to control the odor presentation and were allowed to smell the odor linked to each instruction multiple times. To solidify this learning, they were then tested in a three-alternative forced-choice test. After smelling each odor, they indicated if it was linked to the remember instruction, the forget instruction, or neither instruction. This test ended only after they chose the correct choice for both odors (they were all presented again if there was an incorrect response).

Each trial of the Directed-Forgetting Task included a single target word that was either to-be-remembered or to-be-forgotten. The set of 108 words was randomly divided into practice words (6), buffer words (6), target words (48), and new words, which were only presented in the post-nap test (48). Each trial started with a countdown to zero.

The countdown began with the numbers “4” and “3” (presented on screen for 1 s each). In place of “2”, participants were shown a target word (e.g., “GOLF”) for 1 s, followed by “1” for 1 s. Next, the word “SNIFF” appeared on the screen for 2 s (Fig 1c). The odor presentation was timed so that it would be perceivable at the onset of the “SNIFF” instruction and remain available for ∼2 s. Next, an inter-trial interval of 15 s commenced, during which participants viewed a central fixation cross, and unscented air was presented. The 48 encoding words were shown in three blocks (16 words per block).

Additionally, three buffer words appeared at the beginning and three at the end of each block to avoid primacy and recency effects. In total, each block consisted of 22 words. Before the first block, participants completed a practice round with six target words, half paired with the Remember Odor and half with the Forget Odor, prior to starting the main task. At the end of the practice round, they were asked to type the target words that were linked to the remember instructions and were asked not to type in target words that they were instructed to avoid learning. They then continued to the main task. After each of the three blocks, participants were encouraged to remove their odor mask and walk outside of the room for roughly 1–2 min to reduce odor habituation.

At the conclusion of the Directed Forgetting Task, the Odor-matching Task was administered again with the Remember, Forget, and Unassigned odors (4 s each). Most participants matched each odor with the correct instruction, but two participants did not, and their data were excluded from analysis.

### Spatial Memory Task

Participants removed their odor mask for the Spatial Memory Task (Fig 1d). This task consisted of five blocks, each with 12 objects (randomly selected for each subject). A practice block with four objects was given before starting the first block. Each block consisted of a learning phase and a training phase. During the learning phase, each object was presented in its position on a gray background for 2 s along with the congruent sound, which was played twice, once when the object first appeared and a second time co-terminating with the disappearance of the object. This was followed by an ISI of 2 s that included a fixation cross. Each object was positioned in a unique location with a minimum difference of 120 pixels between objects. After the learning phase, participants began the training phase.

In each self-paced training trial, an object was presented in the center of the screen along with the sound, and the participant had to drag it to its veridical location using the mouse. After they dragged and dropped the item, two images of the object were presented in the placed and veridical location. If the distance between the two was <180 pixels, the placement was considered correct, and the sound was played again. If the placement was incorrect, a line was displayed connecting the object in the placed and veridical location, and the sound was presented. The feedback was presented for 3 s, followed by an inter-trial interval of 2 s, during which a fixation cross was presented. The training phase continued until participants placed each object in its veridical location on two consecutive trials (i.e., the learning criterion was reached). Once the participants reached criterion for any individual object, it was dropped from training, and training continued for the remaining objects. Across participants, a mean of 2.91 ± 0.11 trials were required for an object to reach criterion. Within each block, participants learned and trained to criterion. The order of the objects presented during the practice block and experimental blocks was randomized.

### Pre-Sleep Test

After learning all 60 objects, participants were tested on all of them without feedback. During the test, each object was presented at the center accompanied by its sound, and participants had to drag it to its veridical location. After they placed the object, the sound was played again, followed by a fixation cross for 2 s.

No feedback was provided, and the test trials were self-paced. The first four trials included the objects from the practice block.

### Nap

Participants were offered a 90-min nap opportunity after being fitted with a 32-channel cap with active Ag/AgCl EEG electrodes, as well as two mastoid electrodes, two electrodes placed next to the eyes (EOG), an electrode on the chin (EMG), and the odor mask. EEG scalp locations followed the standard 10–20 layout. Polysomnographic data were recorded using a BrainVision system with a 1000-Hz sampling rate. Data were referenced to the average of the mastoids and bandpass-filtered at 0.5–35 Hz.

Throughout the nap, the lights were turned off, fresh air was flowing continuously through the mask, and white noise was presented through speakers placed approximately 150 cm from the bed. When participants entered stable SWS, odors and sounds were presented unobtrusively. However, if participants did not enter stable SWS sleep in the first 45 mins of the nap, cues were presented during stable N2 sleep (Fig 1e).

The 60 object sounds presented during the spatial memory task were divided into four sets of 15 sounds. Set allocation was determined based on performance on the Pre-Sleep Test (i.e., the between-set variance of the average differences in pre-sleep error was minimized). One set was designated to be presented along with the Remember odor, another set with the Forget odor, another set with the Unassigned odor, and the last set was not presented during sleep (Non-cued). Due to a programming error, a small number of sounds (1.57 ± 0.58) assigned to the first three sets were not presented during sleep and therefore were considered part of the Non-cued set in all analyses.

Stimulus presentation during sleep was divided into blocks, with each block consisting of a single odor and five sounds. In each block of cueing, the odor was presented for 24 s without regard to respiration phase, such that odor inhalation and corresponding sensory processing happened at idiosyncratic times during this 24-s period. With a typical rate of 12-20 respiration cycles per minute, there would have been about 5-8 inhalations while the odor was available. The first sound was presented 3 s after odor onset. Subsequently, the other four sounds were played with a 4.5-s onset-to-onset interval. The odor remained available for 3 s after the last sound’s onset. Each block was followed by a 20-s odorless inter-block interval. Nine such blocks (three conditions x three blocks x five sounds) made up one full round of cueing, which lasted ∼6.27 min. The order of blocks within a round was randomized, and so was the order of sounds within a block.

Cueing was controlled manually by a skilled experimenter and stopped at any sign of arousal, awakening, or transition to REM sleep. After a stop, cueing resumed with the next stimulus after the participant reentered stable NREM sleep. For instance, if an arousal occurred after the second sound in the second block, the odor presentation would stop immediately. Upon resuming, the next cueing block would be shorter and include the presentation of the odor, followed 3 s later by the third, fourth, and fifth sounds of the originally initiated block. If cueing stopped and resumed within 20 s (the length of the inter-block odorless period), fresh air was circulated for 20 s before odor presentation was resumed. On average across participants, 3.13 ± 0.71 blocks (16.90 ± 4.60%) of cueing were interrupted, whereas the remainder included all five sounds. We required at least one round of cueing (45 sounds) for a participant’s data to be included in the analyses.

### Post-Sleep Tests

After waking up, participants took 5-15 min to clean up and then completed two memory tests (Fig 1f). The first test was on the 60 object locations, identical in format to the Pre-Sleep Test. The second test was a recognition test on words from the Directed Forgetting Task. In each self-paced trial, each word appeared along with the question, “Did this word appear in the previous list?” This test included 48 old words (half to-be-remembered and half to-be-forgotten) and 48 new words, presented in a random order. Response choices were “old” or “new”. Participants were instructed to respond regardless of the odor-based instructions. That is, they were told to endorse old words even if they had been presented with the Forget odor. Note that we tested recognition only after the nap, but not before it. This design choice was made intentionally, as instructions to retrieve the to-be-forgotten words could have potentially recontextualised the odors prior to sleep, compromising our main manipulation.

After the Post-Sleep Tests, participants were debriefed and asked if they remembered hearing or smelling anything during sleep. None of the participants recalled hearing or smelling any specific sound or odor.

### Statistical analyses

#### Electroencephalography

EEG preprocessing was conducted using the FieldTrip toolbox (Oostenveld et al., 2011). EEG artifacts were manually detected by an experimenter for rejection. Noisy electrodes were interpolated based on neighboring electrodes using a triangulation method. Sleep scoring was conducted online while the participant was sleeping to determine when to initiate or terminate cueing. Offline sleep scoring was conducted by three independent scorers using the sleepSMG package (https://sleepsmg.sourceforge.net/). Both online and offline scoring were based on the guidelines of the American Academy of Sleep Medicine (Berry et al., 2017). Scorers were not privy to when cueing was done. Any differences in scoring were reconciled among all three scorers.

#### Word recognition test

The results from the word recognition test (Fig 1f) were considered as a manipulation check, confirming that the odors had instruction-specific effects on the preceding words in the Directed Forgetting Test. The percentage of words correctly identified as old was calculated for the Remember and Forget conditions. We ran a paired *t*-test to test the effects of word condition (Remember × Forget) on the percentage of words identified as old using the *ttest* function on MATLAB R2022b. Data was visualized using R Statistical software (v. 4.2.2; R Core Team 2021). Descriptive statistics are provided as mean ± SE.

#### Spatial memory results

Analyses included paired *t*-tests, ANOVAs, and linear mixed models. For the spatial memory task, the error in pixels was calculated as the Euclidean distance between the veridical and participant-placed location. A linear model was used to test the effects of odor conditions on pre-sleep error [Pre-sleep error ∼ Odor condition + (1|Participant)]. The change in memory was calculated as the difference between the error in the pre-sleep and post-sleep test (i.e., positive values indicate memory improvement). We used a linear mixed model to test the effects of odor conditions on change in memory [Change in memory ∼ Odor condition + (1|Participant)]. We ran a *t*-test to compare the change in memory collapsed across the three odor conditions (i.e., the cued conditions) relative to the Non-cued condition. An additional linear mixed model used the change in memory as the dependent variable with the pre-sleep error and odor conditions as the fixed variables [Change in memory ∼ Odor condition * Pre-sleep error + (1|Participant)]. A linear model was used to compare change in memory as a function of the pre-sleep error between the cued conditions (collapsed over three conditions) and the Non-cued condition [Change in memory ∼ Cueing condition * Pre-sleep error + (1|Participant)]. The *anova* function for linear mixed-effects models in MATLAB was used to test the significance of the linear models. For data analysis and visualization, we used MATLAB R2022b (MathWorks Inc., Massachusetts, USA) and R Statistical software (v. 4.2.2; R Core Team 2021), respectively.

#### Spectral analysis

Continuous EEG data were segmented relative to odor and sound onsets during sleep. Data were segmented in two different ways: with epochs starting at 10 s before and ending 45 s after odor onset, and with epochs starting at 1.5 s before and ending 3 s after sound onset. Spectrograms were computed using MATLAB R2022b with the following parameters: 0.5 Hz to 20 Hz with 0.5-Hz increments; 500-ms sliding window with 80% overlap. Spectrograms were normalized by considering the percentage change in power relative to a baseline period (5 s before odor onset and 0.5 s before sound onset) as done in prior studies (e.g., Schechtman et al., 2021). Analyses were conducted on data averaged across a cluster of seven central electrodes: two frontocentral, three central electrodes including Cz, and two centroparietal electrodes (FC1, FC2, C3, Cz, C4, CP1, and CP2). As done in previous studies (e.g, Schechtman et al., 2021; Yao et al., 2024), electrodes around the midline were chosen as their activity reflects sleep spindles and slow-wave activity. Only data collected during NREM sleep were included in the analyses, thus avoiding arousal effects (see Fig S1 for analysis of data from all trials, regardless of sleep stage). As mentioned above, if cueing was interrupted mid-block, the remainder of the block was presented the next time the participant entered stable NREM sleep. As a result, some of these partial blocks were shorter than others. Both partial and full blocks were included in analyses, as long as no arousal or transition to another sleep stage was observed in the data (i.e., as long as the partial or full cueing period was executed during NREM).

To identify significant clusters of activity in response to sound onset, we averaged data across segments encompassing all sounds for each participant and ran *t*-tests across participants for each point in time-frequency space (corrected for multiple comparisons using Family-wise corrections). Two significant clusters in the sigma (12–16 Hz) and delta-theta (2–8 Hz) range were identified, and we calculated the change in power per trial within these significant clusters for each trial. We then ran two linear mixed models using the power change in the clusters for sigma and delta-theta band as the dependent variable and the odor condition as the independent variable.

Model 1: Power in sigma range ∼ Odor condition + (1|Participant) Model 2: Power in delta-theta range ∼ Odor condition + (1|Participant) The *ANOVA* function for linear mixed-effects models in MATLAB R2022b was used to test the significance of both linear models.

Lastly, to examine the change in the power in the sigma (12–16 Hz), delta (0.5–4 Hz), and alpha (8–11 Hz ) ranges, we averaged power in 5-s epochs beginning 5 s before and ending 45 s after odor onset. Power was baseline-corrected for the 5 s before onset. Considering the absence of an Odor condition effect in prior linear models, the analysis on change in power data was conducted after collapsing across odor conditions. We conducted one-sample *t*-tests comparing data in each epoch to zero to determine significant power changes. For each frequency band, we corrected for multiple comparisons using family-wise corrections.

## Results

### Odor cues influenced memory for words

During the Directed Forgetting Task, participants were presented with words, followed by odors instructing them to either remember (24 words) or forget the preceding word (24 words). Recognition memory for the words was tested after the afternoon nap. This is a precondition for the success of our approach – if odors were not sufficiently linked to the instructions during wakefulness, they would be unlikely to have an impact on memory during sleep. As shown in Fig 2, memory was better for Remember words (86.24 ± 3.05%) than for Forget words (55.43 ± 3.05%). A paired *t*-test revealed a significant difference [*t* (22) = 5.06, *p* < .001]. The mean false alarm rate was 11.24 ± 2.11% for new words. These results confirm that participants successfully linked instructions to the respective odors and that we were able to replicate the directed-forgetting effect using odor cues.

**Figure 2.**
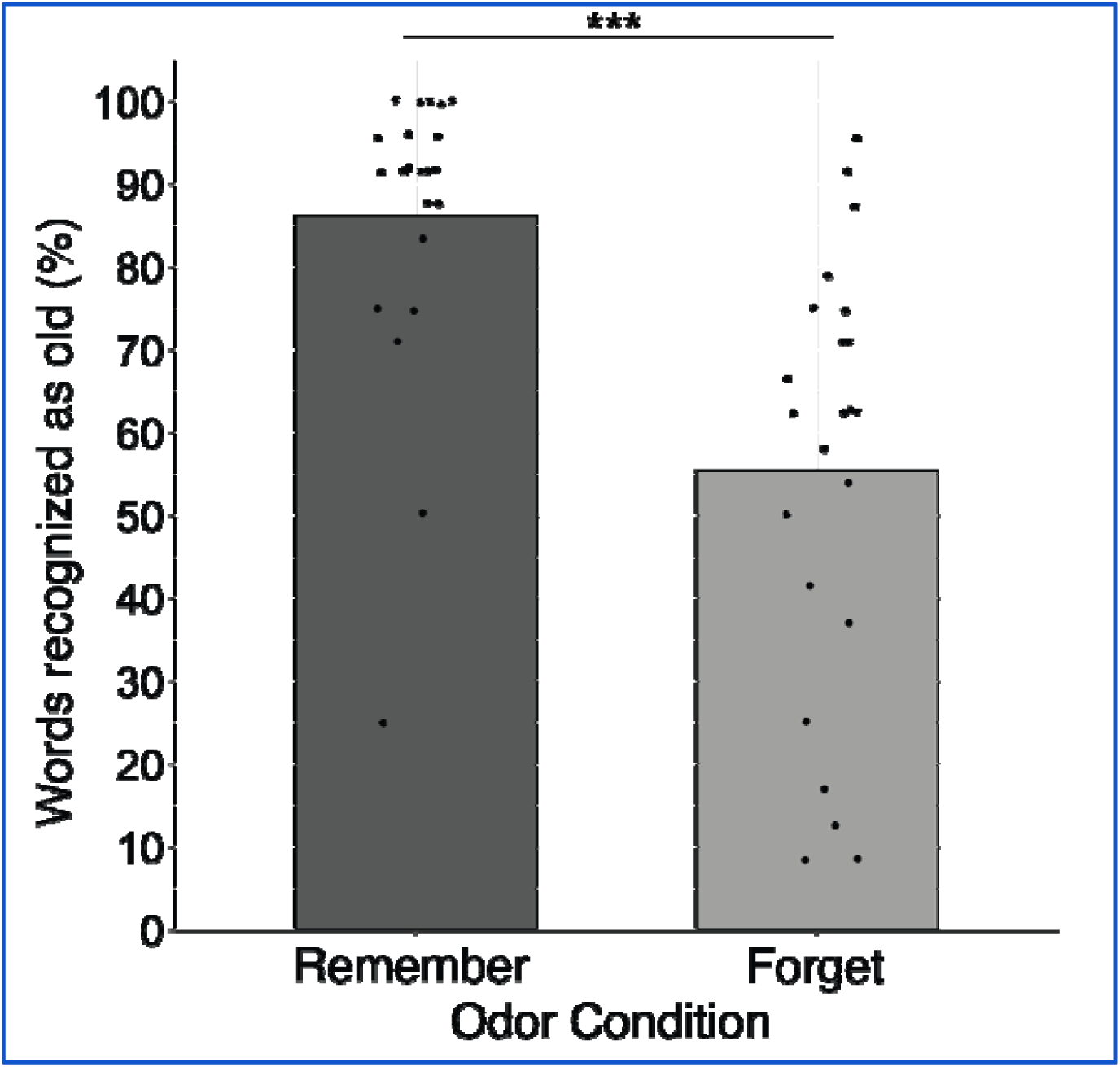
Results from the Directed Forgetting Task. Remember words were recognized more than Forget words, indicating that the odors were effective in conveying directed-forgetting instructions. Each dot represents an individual participant. *** *p* < .001

### Sound cues improved memory most when presented with the unassigned odor

Participants completed a Spatial Memory Task wherein each object was associated with a unique on-screen location and a congruent sound. The spatial recall was tested before and after an afternoon nap. During the nap, object sounds were divided into four sets. Two sets were paired with the Remember odor and Forget odor from the Directed Forgetting Task, a third set with an Unassigned odor and a fourth set was not presented during sleep. The mean pre-sleep error was 184.43 ± 10.07 pixels (4.88 ± 0.27 cm) and did not differ across the four conditions [Remember, Forget, Unassigned, and Non-cued; *F* (3,1376) = 0.20, *p* = .89]. The mean post-sleep error was 204.95 ± 12.75 pixels (5.42 ± 0.34 cm). The pre/post change in recall accuracy is shown in Fig 3a. A linear model considering the effects of condition on change in memory from pre-sleep to post-sleep failed to reveal a significant effect [*F* (3,1376) = 0.72, *p* = .54]. Contrary to our prediction, the odors associated with Remember or Forget instructions did not influence the change in memory for the co-activated spatial memories cued by the sounds. Furthermore, change in memory was not significantly different when comparing between objects that were reactivated during sleep (collapsed over the three cued conditions) and those that were not (the Non-cued condition; *t* (22) = 0.59, *p* = .56), indicating an absence of a classic TMR memory benefit observed in prior studies (Hu et al., 2020).

**Figure 3.**
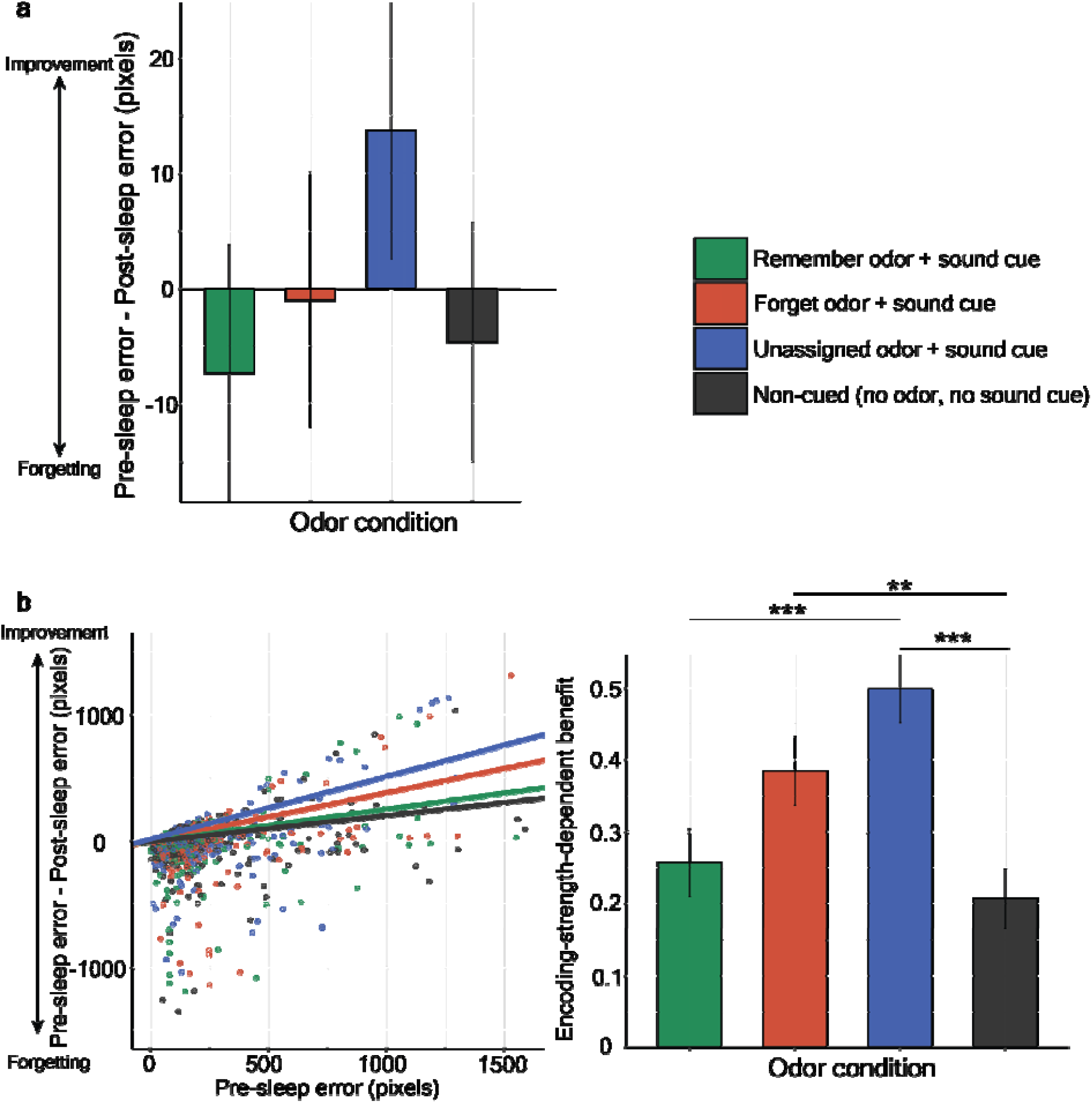
Sleep reactivation impacted memory most for object locations paired with the Unassigned odor in a memory-strength-dependent manner. **a** The change in memory across all items was visualised using the coefficients of the linear mixed model accounting for the random effects of participants. The change in memory (in pixels) from pre-sleep to post-sleep was not significantly different across conditions. **b (left)** Change in memory from pre-sleep to post-sleep as a function of pre-sleep error for all four conditions. Each dot represents one of the 60 objects per participant (but note that the model included random effects for participants that are not reflected here). The right tail of the graph reflects objects with high pre-sleep error (i.e., poorly encoded). Absolute memory benefits would be reflected by a difference in the intercept between conditions; Steeper slopes indicate larger memory benefits for poorly encoded objects. reactivation improved memory for poorly encoded objects when paired with the Unassigned odor (and vice versa for strongly encoded memories). **b (right)** The slopes of the curves presented in panel (b) showed significant differences between the Unassigned condition and both the Non-cued and Remember conditions. Higher values indicate more encoding-strength-dependent benefits. Error bars indicate SEM adjusted for within-subjects comparisons. * *p* < .05, *** p < .001

Another linear model considering the effects of pre-sleep error and odor condition on change in memory revealed a significant pre-sleep effect on change in memory [*F* (1, 1372) = 832.4, *p* < .001] and a lack of odor effect on change in memory [*F* (3, 1372) = 0.70, *p* = .55]. However, the interaction between pre-sleep error and condition indicated that initial memory strength modulated how odors impacted memory [*F* (3,1372) = 0.46, *p* <.001]. That is, the relationship between pre-sleep error and the benefit following sleep varied with condition, as shown in Fig 3b, left and right. Higher slopes reflect larger benefits for weakly encoded memories (and vice versa for strongly encoded memories), and results for objects presented with the Unassigned odor demonstrated the steepest slope (see Table 1 for pairwise comparisons). The slope of the Unassigned odor condition was significantly larger than the Remember and Non-cued conditions, and marginally larger than the Forget odor condition. Surprisingly, the slope for the Forget condition was higher than for the Non-Cued condition, suggesting that, contrary to our predictions, the Forget condition benefited rather than weakened memory (at least for weakly encoded objects). Further, the encoding-strength-dependent TMR benefit was significantly larger for the objects that were reactivated during sleep (i.e., collapsed across all odor conditions) compared to the Non-cued condition [*F* (1,1376) = 12.22, *p* < .001].

**Table 1.** Pairwise comparisons for the interaction between pre-sleep error and condition *The t-statistic, degrees of freedom, and p-values for the pairwise comparisons of the slope in the linear mixed model examining the effects of conditions and pre-sleep error on change in memory. Positive t-statistics indicate that Condition 1 > Condition 2 in terms of slope*.

| **<u>Condition 1</u>** | **<u>Condition 2</u>** | **<u>t statistic (df)</u>** | **<u>p-value</u>** |
| --- | --- | --- | --- |
| **Unassigned** | **Non-cued** | 4.66 (1372) | <.001*** |
| **Unassigned** | **Remember** | 3.62 (1372) | <.001*** |
| **Unassigned** | **Forget** | 1.70 (1372) | 0.09 |
| **Non-cued** | **Remember** | -0.80 (1372) | 0.42 |
| **Non-cued** | **Forget** | -2.81 (1372) | 0.005** |
| **Remember** | **Forget** | -1.89 (1372) | 0.06 |

\* $p < .05$ , \*\* $p < .01$ , \*\*\* $p < .001$

### Odor cues were followed by a prolonged decrease in sigma power

During the 90-min nap opportunity, each of the three odors was paired with 15 sounds. An additional set of 15 sounds was not presented during sleep. Only cues presented during NREM stages (N2 and SWS) were included in the analyses (*n* = 908, 898, 896 sound cues for the Remember, Forget, and Unassigned odor conditions, respectively; trials for each condition were grouped across participants). Only cues presented during NREM stages were analyzed to avoid any effects of arousal. Table 2 shows details on sleep characteristics along with the number of sound and odor cues presented. On average across participants, 2.64 ± 0.20 complete rounds of cues were presented. Sleep spindles, waxing and waning waveforms at the sigma frequency band (12–16 Hz), are associated with memory processing and often observed in response to stimuli presented during N2 and SWS sleep (Cairney et al., 2018; Fernandez & Lüthi, 2020). Memory consolidation has also been associated with K-complexes and slow waves (0.5–2 Hz) that are encompassed within the delta (0.5–4 Hz) and theta (4–8 Hz) frequency bands and are frequently observed following cue presentation (Walker, 2009; Xia et al., 2023). Considering the role of spectral responses within these frequency bands on memory processing and the encoding-dependent memory benefit found in the current study, we computed EEG power changes in these frequency bands following sound cues. Although changes in spectral power are not synonymous with spindles, K-complexes, and slow waves, they do reflect these waveforms.

**Table 2.**
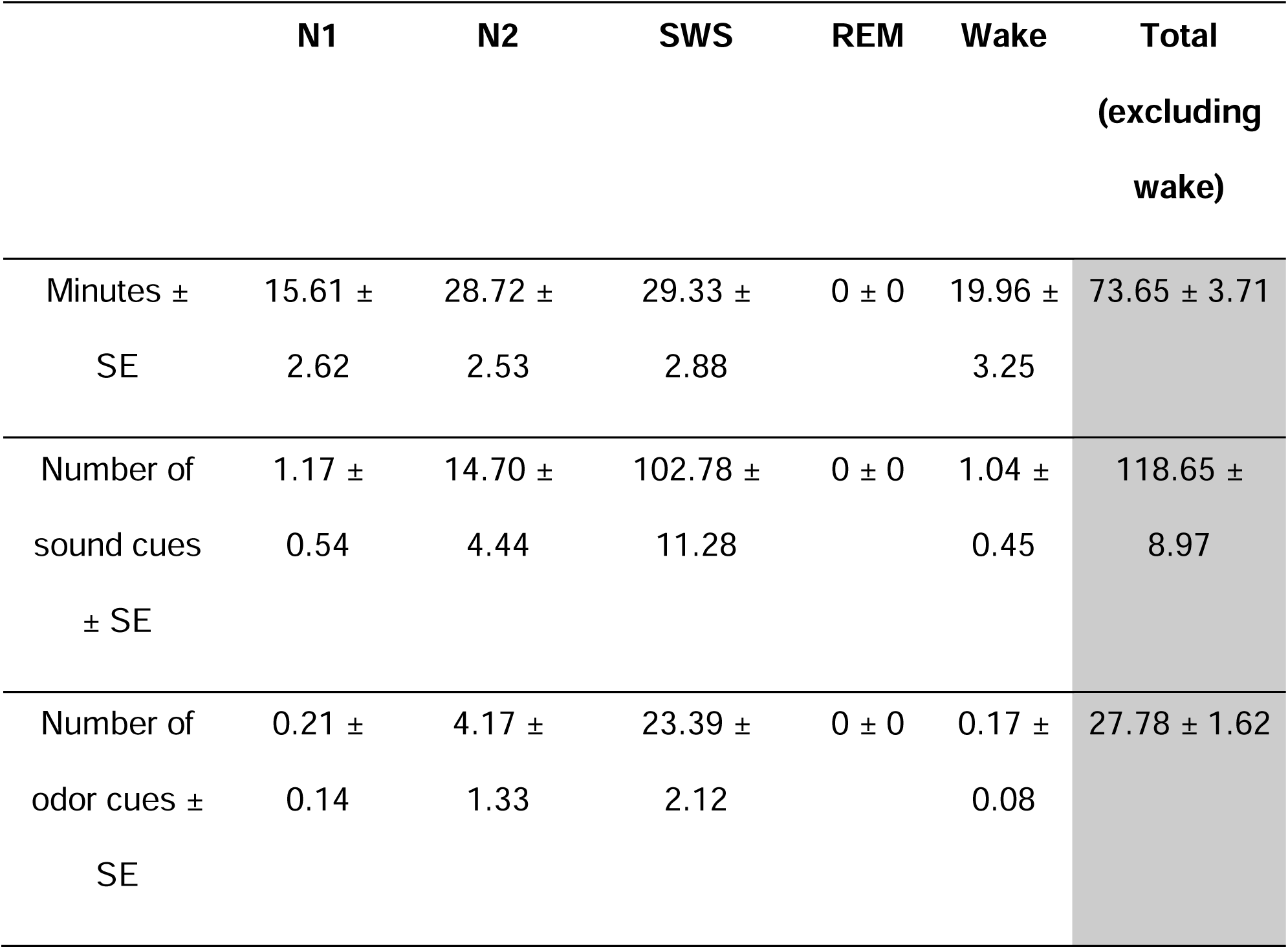
Sleep characteristics and number of cues *The distribution of sleep stages during the nap and mean number of cues in each stage. N1 - stage 1 of sleep, N2 - stage 2 of sleep, SWS - slow-wave sleep, REM - rapid eye movement sleep, SE - standard error*.

Fig 4 shows EEG data as a function of odor condition across the entire 24-s period of odor plus sound presentation (left), as well as time-locked to sound presentations (right). We first used the data collapsed across all sound cues, conditions, and participants to identify clusters of activity. We found significant clusters in the sigma and delta-theta bands after correcting for multiple comparisons ( < .05). Fig S2 illustrates the significant clusters in the sigma (∼15–18 Hz) and delta-theta bands (∼3–9 Hz) identified after correcting for multiple comparisons. We then computed power change in these clusters separately as a function of condition and ran two linear mixed models. In the first model, we evaluated the influence of condition on sigma power change. The model revealed no effect of condition [*F* (2, 2699) = 1.01, *p* = .36]. The second model focused on delta-theta power and similarly revealed no condition effect [*F* (2, 2699) = 2.48, *p* = .08].

**Figure 4.**
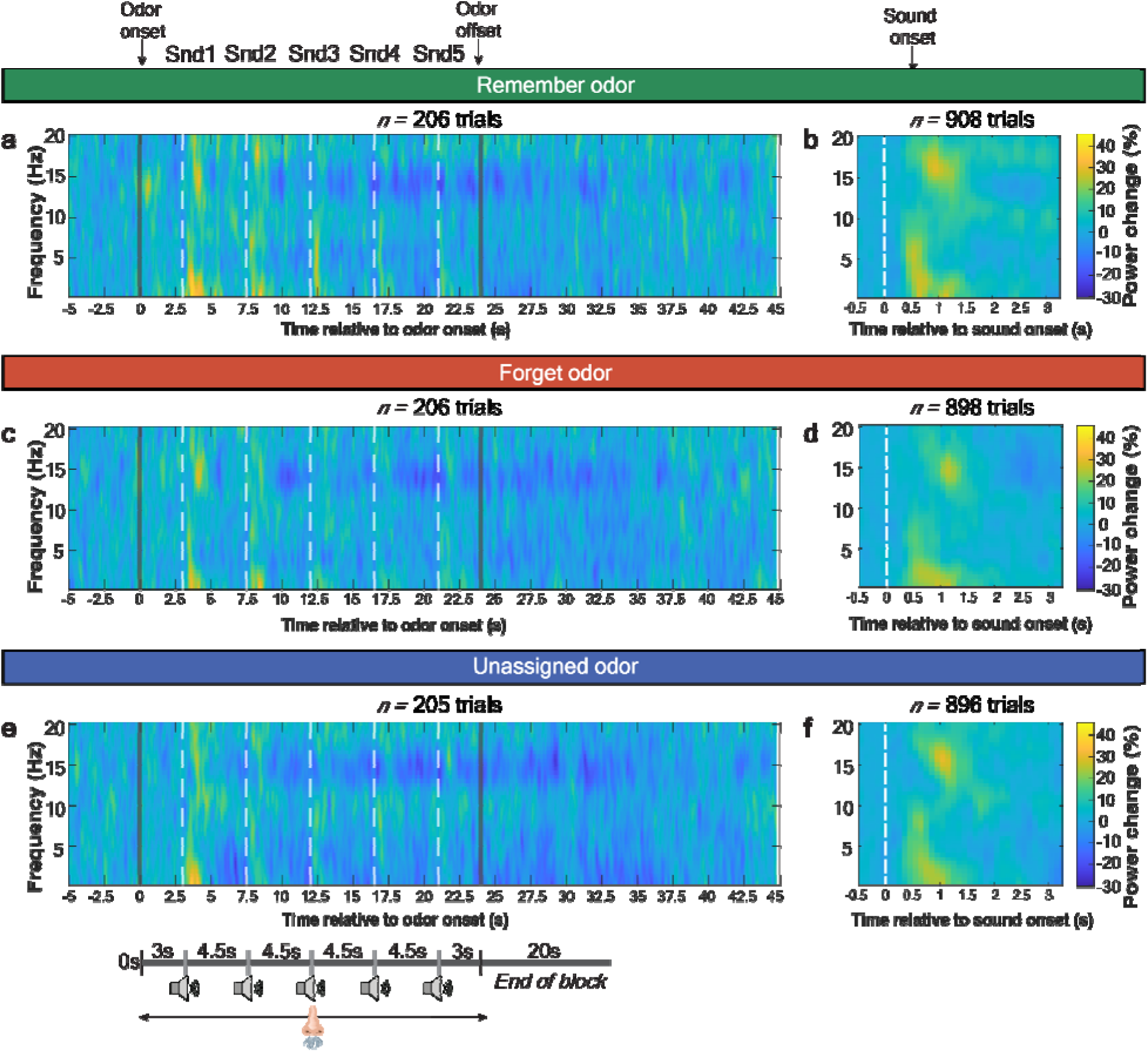
EEG response to cueing did not vary as a function of odor condition. Baseline-corrected spectrograms displaying the power change as a percentage relative to baseline across the frequency bands (0–20 Hz) collapsed across participants. Only trials that commenced and ended within NREM, and therefore did not include arousals, were included in the analysis (this includes partial cueing blocks, consisting of less than five sounds). Data for all trials, regardless of sleep stage, are presented in Figure S1. Gray lines indicate odor onset and offset. Snd1-Snd5 describes the onset of the five sounds presented with a 4.5-s onset-to-onset interval indicated by the white, dashed lines. The panels on the left column (**a,c,e**) describe activity in response to odor onset and the panels on the right column (**b,d,f**) describe spectral activity in response to sound onset.

Sigma and delta-theta activity were also analyzed over the period of odor presentation. We included data from 5 s before to 45 s after odor onset, and included trials only when both the odor onset and offset were during NREM sleep (n= 206, 206, 205 odor presentations for the Remember, Forget, and Unassigned odor conditions, respectively). Analyses were collapsed over conditions. Results demonstrated decreases in power in the sigma band. In a post-hoc analysis, we analyzed data in 5-s epochs, focusing on average power for the sigma (12–16 Hz), delta (0.5–4 Hz), and alpha (8–11 Hz) bands, as shown in Fig 5. For each band, we corrected for multiple comparisons using family-wise corrections. No significant variation was observed in alpha and delta, whereas sigma power increased between 0–5 s after odor onset and declined for all epochs between 10–35 s after odor onset (*p* < .001, corrected). The initial increase in sigma power could be attributed to an increase in spindle activity in response to odor or sound onset, as observed in prior studies (Rihm et al., 2014; Schechtman et al., 2021). However, the long-lasting decline (25 s, ending more than 10 s after odor offset) is unlikely to reflect a post-spindle refractory period, which is usually limited to 4–6 s (Antony et al., 2018). The probability of spindle occurrence exhibited a similar decline from 15 s to 35 s after odor onset (Fig S3). This finding indicated that the sigma decline observed corresponds to a decline in spindle activity in response to odor presentation.

**Figure 5.**
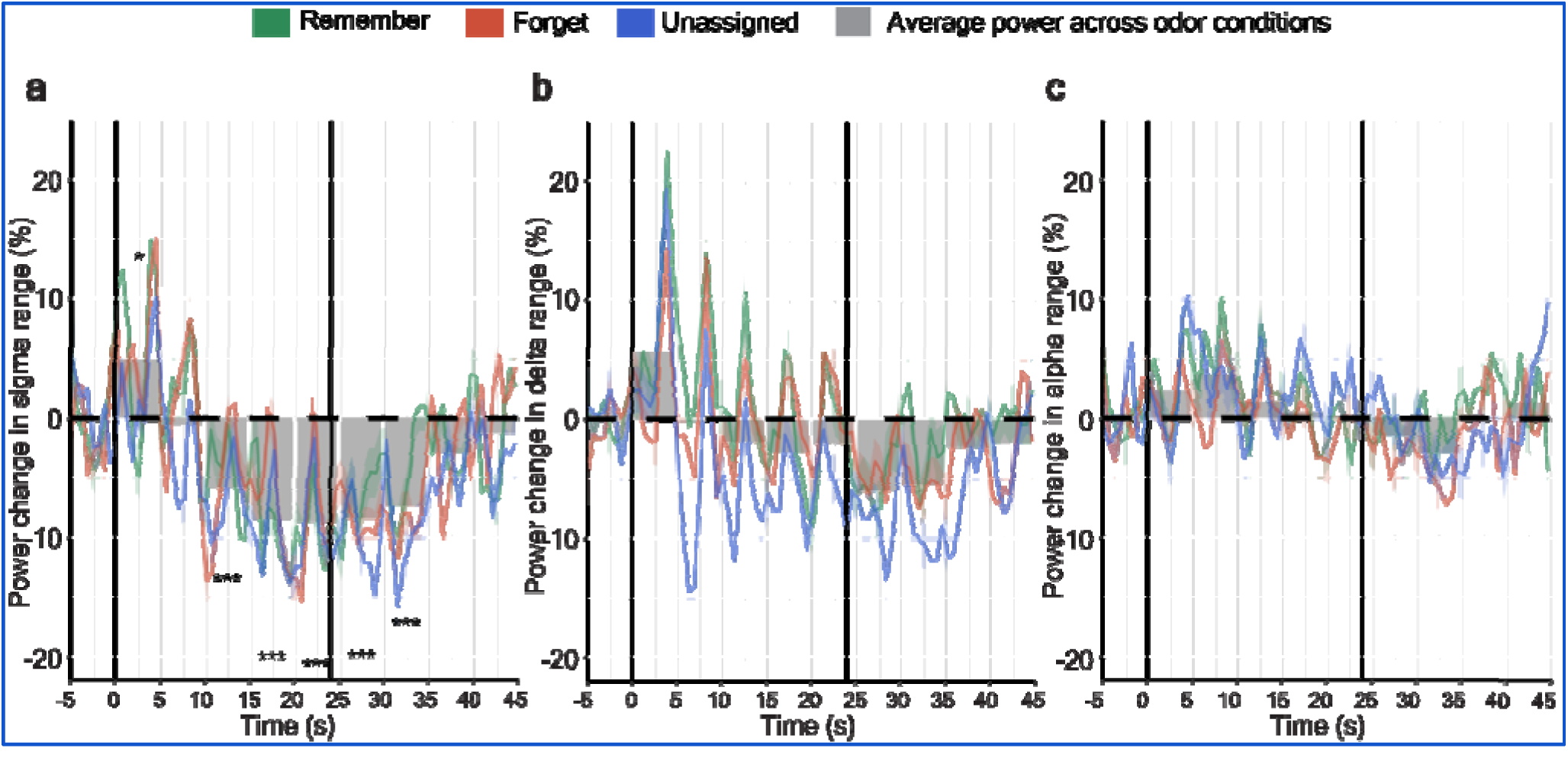
Odor cues were followed by a prolonged decrease in sigma power. **a** Baseline-corrected power change (%) in the sigma band (12–16 Hz) during odor and sound presentation from 5 s before to 45 s after odor onset. The gray bars represent the averaged power change across conditions computed for 5-s epochs from 5 s before to 45 s after odor onset (i.e., 10 epochs total). Sigma power initially increased between 0 s and 5 s, which could be attributed to an increase in spindle activity in response to odors or subsequent sounds. Then, sigma power declined significantly from 10 s to 35 s after odor onset. Interestingly, the power decline persisted after odor offset. **b** Similarly, baseline-corrected power change (%) in the delta (0.5–4 Hz) was plotted as a function of time (s) relative to odor onset (at 0). The power did not vary significantly after odor onset. **c** Lastly, variations in the power change (%) in the alpha band (8–11 Hz) were plotted against time (s). Alpha power did not vary significantly after odor onset. Black vertical lines indicate odor onset and offset. *p*-values were corrected for multiple comparisons. \**p* < .05, \**\*p* < .01, \*\*\**p* < .001

## Discussion

The current study aimed to modulate memory consolidation by simultaneously presenting odors and sounds during sleep. Odors were first introduced during wakefulness to instruct participants to either Remember and Forget words, with the expected effects on memory, better recognition for Remember compared to Forget words. We hypothesized that these two odors, presented during sleep, could reactivate brain states involved in motivated remembering or forgetting, and thereby impact memories. To the contrary, we did not observe accuracy differences for memories reactivated in the presence of the Remember odor versus the Forget odor. Memory accuracy was also similar for memories reactivated in the presence of the Unassigned odor (i.e., no assigned meaning before sleep) and memories not reactivated at all.

A further analysis showed that reactivation did impact memory in an encoding-strength-dependent manner, as seen in other TMR studies (e.g., Cairney et al., 2016; Lazarus et al., 2025; Schechtman et al., 2023). That is, memory accuracy was improved to the extent that larger errors were observed on the Pre-Sleep Test (i.e., for weakly encoded memories). Of note, the term weakly encoded is based on spatial accuracy rather than the retention of memories over time. These findings complement others demonstrating that TMR is most effective for weakly encoded memories. For example, Schechtman and colleagues (2023) showed that TMR impacted memory in an encoding-strength-dependent manner not only for cued memories, but also for their contextually related counterparts. Creery and colleagues (2015) analyzed results for individual items learned in a spatial memory task and reported that sound-cued reactivation produced a larger benefit for those items with a higher pre-sleep error compared to those with less pre-sleep error. Similarly, Cairney and colleagues (2016) showed that TMR improved memory selectively for picture locations that demonstrated low retrieval accuracy before sleep. These findings from TMR studies complement other studies showing that overnight sleep prioritizes weaker memories, benefitting memories based on their strength before sleep (e.g., Petzka et al., 2021; Wernette & Fenn, 2024).

We predicted that spatial accuracy would be enhanced when object sounds were presented concurrently with the Remember odor and the reverse with the Forget odor. Contrary to this prediction, odors did not reliably influence change in memory accuracy . The observed encoding-strength-dependent benefit was not significantly different for the Non-cued and Remember conditions. The encoding-strength-dependent effect was significantly larger for the Forget condition when compared to the Non-cued condition, indicating a lack of a suppression effect. Interestingly, the largest encoding-strength-dependent benefit was observed for the Unassigned condition.

One interpretation of these unpredicted results is that odors that had a meaningful link before sleep – the Remember and Forget odors – may have nullified the memory benefits that would have otherwise been produced by the sound cues.

Previous studies have shown that auditory cues presented in close succession during sleep disrupt reactivation (Farthouat et al., 2017; Schreiner et al., 2015). Here, odors were continually present during sound presentations, and given that odors are taken in with each inhalation, odor processing likely occurred close in time to the auditory cues. The motivation behind our study was to test whether presenting reactivation cues from different modalities might facilitate concurrent reactivation without interference. Our results suggest that sounds did not effectively reactivate object memories when presented along with an odor stimulus from the unrelated Directed-Forgetting task.

Perhaps odor cues reactivated the instructions to remember or forget sufficiently but this reactivation could have interfered with the spatial memories brought forth by the sound cues. Such interference may explain the forgetting effect observed by Simon and colleagues (2018) when they paired a Forget tone with sound cues from an unrelated spatial task. This interpretation is in line with a recent hypothesis that concurrent memory reactivation only benefits subsequent retrieval if said memories are tightly integrated and complementary (Antony & Schechtman, 2023). If correct, this idea highlights a limitation of sleep-related reactivation: only tightly interlinked memories can be reactivated together without interference.

A related idea is that the Remember and Forget odors produced reactivation of the various episodic features associated with these odors before sleep. Both odors were presented repeatedly during the Directed-Forgetting Task (33 times each). Each trial constituted an episode including a countdown, a word presentation, and a sniff instruction, such that one odor or the other was perceived and its assigned meaning activated. Odor presentations during sleep may have reactivated some of this episodic information instead of or in addition to the Remember or Forget instruction. Less interfering memory reactivation was likely produced by the unassigned odor, given the absence of preassigned meaning and associated learning episodes, such that sound cues could benefit spatial memory. We also note that our cueing regimen led to a relatively high level of arousals and may have compromised the efficacy of TMR to a certain degree, as reported in other studies (Göldi & Rasch, 2019; Whitmore & Paller, 2023).

The results nevertheless expanded the directed-forgetting literature. Typically in such studies, participants are directed to forget (or remember) recently encoded information based on visual (e.g., Scotti & Maxcey, 2022) or auditory cues (e.g., Fawcett & Taylor, 2008). Here, we demonstrated that odors can also be used to effectively convey Directed-Forgetting instructions.

The odors and sounds presented during sleep elicited neural activity across conditions. Based on data collapsed across sounds and the three odor conditions, we identified clusters of EEG activity in sigma (12–16 Hz) and delta-theta (0.5–8 Hz) frequency bands that increased systematically following stimulation. These two bands can be sensitive to sleep spindles and slow waves, respectively, that are characteristic of NREM sleep (Fernandez & Lüthi, 2020; Xia et al., 2023), although power changes in these frequency bands are not limited to these EEG waveforms. A significant increase in sigma activity was observed between 0–5 s after odor onset. It is likely that, in all three conditions, spindles were provoked by individual sound presentations leading to the increase in sigma power. Alternatively, increases in spindle activity could have been in relation to odor cues, as seen previously (e.g., Rihm et al., 2014; Sánchez-Corzo et al., 2024).

In the current study, the temporal dynamics of sigma exhibited a unique pattern: a decline in power during odor presentation beginning roughly 10 s after odor onset and continuing for 35 s (11 s after odor offset). Parallel changes were found in spindle counts, supporting the inference that the sigma decrease reflected changes in spindles. However, the cause for this decrease is unclear. Previous studies have revealed a refractory period following individual spindles that lasts 2–4 s (Antony et al., 2018).

However, the observed decrease is on a different timescale. Another study from our lab similarly revealed a decline in sigma power, which lasted 15 s after odor offset during sleep (Narayan et al., [unpublished]). Recently, studies have shown that spindle densities fluctuate at an infra-slow rate of ∼0.02 Hz (Fernandez & Lüthi, 2020; Lecci et al., 2017), possibly driven by fluctuations in noradrenaline in the locus coeruleus (Osorio-Forero et al., 2025). The temporal dynamics of the decline in sigma power and spindle probability in our study are of the same order of magnitude as these endogenous fluctuations, suggesting that TMR stimuli might entrain them, although the mechanism through which this may happen remains unclear. Alternatively, the prolonged odor presentation could have caused habituation and altered the breathing patterns. Odorants presented during sleep have been observed to influence respiration, decreasing inhalation and increasing exhalation (Arzi et al., 2010). Such a modulation of respiration may have caused the prolonged decrease in sigma power.

Our study has several limitations. First, the sample size was smaller than that used in other studies in the field. This may be one of the reasons why we were unable to find an overall cueing benefit. The long cueing sequence, along with the requirement that participants sleep with an odor mask, made data collection more challenging than typical TMR studies. Additionally, the co-presentation of cues in different modalities may itself have increased the probability of arousals. These factors also led to more fragmentation of cueing blocks, which may have impacted the efficacy of TMR to some degree. Another limitation is that we were unable to examine the direct neural response following odor exposure. Due to the absence of respiration data, we were unable to time-lock odor onset to inhalations and evaluate variations in neural activity in response to odor onset. Future research should validate the precise timing of sigma decline based on participants’ respiratory cycles. Lastly, our study did not include a follow-up session, and it may be that a larger influence of odors on spatial memory would emerge with longer delays (e.g., 7 days as in Simon et al., 2018).

The present study attempted to combine odor and sound presentation during sleep, introducing a novel experimental design. As far as we know, this is the first TMR study to use more than one stimulation modality. The concurrent presentation of odors and sounds during sleep was intended to suppress or boost previously established memories, based on the experimental condition. However, the findings from our study cast doubt on whether odors and sounds can be combined in this manner. Temporally segregating stimuli, so that odors are not inhaled near the time of sound presentations, may be required to combine memory cues effectively. Alternatively, the sleeping brain may be unable to co-activate unrelated memories without one blocking the consolidation of another. How other factors might influence whether multiple cues can be combined without interference remains an exciting topic for further investigation.

Further understanding of these aspects of sleep-based consolidation is critical for deciphering the mechanisms whereby memory storage is altered during sleep and for capitalizing on the potential benefits of these alterations.

## Acknowledgements

Supported by the US National Institutes of Health (K99-MH122663, R00-MH122663 to ES; DP1-HL179370 to KAP).

## References

1. Antony, J. W., Piloto, L., Wang, M., Pacheco, P., Norman, K. A., & Paller, K. A. (2018). Sleep Spindle Refractoriness Segregates Periods of Memory Reactivation. Current Biology, 28(11), 1736–1743.e4. 10.1016/j.cub.2018.04.020

2. Antony, J. W., & Schechtman, E. (2023). Reap while you sleep: Consolidation of memories differs by how they were sown. Hippocampus, 33(8), 922–935. 10.1002/hipo.23526

3. Arzi, A., Sela, L., Green, A., Givaty, G., Dagan, Y., & Sobel, N. (2010). The Influence of Odorants on Respiratory Patterns in Sleep. Chemical Senses, 35(1), 31–40. 10.1093/chemse/bjp079

4. Berry, R. B., Brooks, R., Gamaldo, C., Harding, S. M., Lloyd, R. M., Quan, S. F., Troester, M. T., & Vaughn, B. V. (2017). AASM Scoring Manual Updates for 2017 (Version 2.4). Journal of Clinical Sleep Medicine, 13(05), 665–666. 10.5664/jcsm.6576

5. Brodeur, M. B., Dionne-Dostie, E., Montreuil, T., & Lepage, M. (2010). The Bank of Standardized Stimuli (BOSS), a New Set of 480 Normative Photos of Objects to Be Used as Visual Stimuli in Cognitive Research. PLoS ONE, 5(5), e10773. 10.1371/journal.pone.0010773

6. Cairney, S. A., Guttesen, A. Á. V., El Marj, N., & Staresina, B. P. (2018). Memory Consolidation Is Linked to Spindle-Mediated Information Processing during Sleep. Current Biology, 28(6), 948–954.e4. 10.1016/j.cub.2018.01.087

7. Cairney, S. A., Lindsay, S., Sobczak, J. M., Paller, K. A., & Gaskell, M. G. (2016). The Benefits of Targeted Memory Reactivation for Consolidation in Sleep are Contingent on Memory Accuracy and Direct Cue-Memory Associations. Sleep, 39(5), 1139–1150. 10.5665/sleep.5772

8. Creery, J. D., Oudiette, D., Antony, J. W., & Paller, K. A. (2015). Targeted Memory Reactivation during Sleep Depends on Prior Learning. Sleep, 38(5), 755–763. 10.5665/sleep.4670

9. Farthouat, J., Gilson, M., & Peigneux, P. (2017). New evidence for the necessity of a silent plastic period during sleep for a memory benefit of targeted memory reactivation. Sleep Spindles & Cortical Up States, 1(1), 14–26. 10.1556/2053.1.2016.002

10. Fawcett, J. M., & Taylor, T. L. (2008). Forgetting is effortful: Evidence from reaction time probes in an item-method directed forgetting task. Memory & Cognition, 36(6), 1168–1181. 10.3758/mc.36.6.1168

11. Fernandez, L. M. J., & Lüthi, A. (2020). Sleep Spindles: Mechanisms and Functions. Physiological Reviews, 100(2), 805–868. 10.1152/physrev.00042.2018

12. Göldi, M., & Rasch, B. (2019). Effects of targeted memory reactivation during sleep at home depend on sleep disturbances and habituation. *Npj Science of Learning*, *4*(1), 5. 10.1038/s41539-019-0044-2

12. Hu, X., Cheng, L. Y., Chiu, M. H., & Paller, K. A. (2020). Promoting memory consolidation during sleep: A meta-analysis of targeted memory reactivation. Psychological Bulletin, 146(3), 218–244. 10.1037/bul0000223

13. Joensen, B. H., Harrington, M. O., Berens, S. C., Cairney, S. A., Gaskell, M. G., & Horner, A. J. (2022). Targeted memory reactivation during sleep can induce forgetting of overlapping memories. Learning & Memory, *29*(11), 401–411. 10.1101/lm.053594.122

14. Lazarus, A., Bassard, A., & Schechtman, E. (2025). Manipulating memory processing during sleep to explore the critical duration of reactivation events. Neuropsychologia, 217, 109211. 10.1016/j.neuropsychologia.2025.109211

15. Lecci, S., Fernandez, L. M. J., Weber, F. D., Cardis, R., Chatton, J.-Y., Born, J., & Lüthi, A. (2017). Coordinated infraslow neural and cardiac oscillations mark fragility and offline periods in mammalian sleep. Science Advances, 3(2). 10.1126/sciadv.1602026

16. Narayan, G., Babineaux III, G., Cho, M., Murugavel, S., Lu, T., Lew, N. J., Aquino Argueta, S., & Schechtman, E. (n.d.). *Examining the Specificity of Memory Reactivation During Sleep using Odor-Cued Targeted Reactivation* [Unpublished].

17. Oostenveld, R., Fries, P., Maris, E., & Schoffelen, J.-M. (2011). FieldTrip: Open Source Software for Advanced Analysis of MEG, EEG, and Invasive Electrophysiological Data. Computational Intelligence and Neuroscience, 2011, 1–9. 10.1155/2011/156869

18. Osorio-Forero, A., Foustoukos, G., Cardis, R., Cherrad, N., Devenoges, C., Fernandez, L. M. J., & Lüthi, A. (2025). Infraslow noradrenergic locus coeruleus activity fluctuations are gatekeepers of the NREM–REM sleep cycle. Nature Neuroscience, 28(1), 84–96. 10.1038/s41593-024-01822-0

19. Oudiette, D., & Paller, K. A. (2013). Upgrading the sleeping brain with targeted memory reactivation. Trends in Cognitive Sciences, 17(3), 142–149. 10.1016/j.tics.2013.01.006

20. Petzka, M., Charest, I., Balanos, G. M., & Staresina, B. P. (2021). Does sleep-dependent consolidation favour weak memories? Cortex, 134, 65–75. 10.1016/j.cortex.2020.10.005

21. Poe, G. R. (2017). Sleep Is for Forgetting. The Journal of Neuroscience: The Official Journal of the Society for Neuroscience, 37(3), 464–473. 10.1523/JNEUROSCI.0820-16.2017

22. Rihm, J. S., Diekelmann, S., Born, J., & Rasch, B. (2014). Reactivating Memories during Sleep by Odors: Odor Specificity and Associated Changes in Sleep Oscillations. Journal of Cognitive Neuroscience, 26(8), 1806–1818. 10.1162/jocn_a_00579

23. Sánchez-Corzo, A., Baum, D. M., Irani, M., Hinrichs, S., Reisenegger, R., Whitaker, G. A., Born, J., Sitaram, R., & Klinzing, J. G. (2024). Odor cueing of declarative memories during sleep enhances coordinated spindles and slow oscillations. NeuroImage, 287, 120521. 10.1016/j.neuroimage.2024.120521

24. Santangelo, V., Pedale, T., Daviddi, S., Salsano, I., Macrì, S., & Campolongo, P. (2025). Altered brain activity during active forgetting in highly superior autobiographical memory: Evidence from an item-method directed forgetting. iScience, 28(6), 112607. 10.1016/j.isci.2025.112607

25. Schechtman, E., Antony, J. W., Lampe, A., Wilson, B. J., Norman, K. A., & Paller, K. A. (2021). Multiple memories can be simultaneously reactivated during sleep as effectively as a single memory. Communications Biology, 4(1), 25. 10.1038/s42003-020-01512-0

26. Schechtman, E., Heilberg, J., & Paller, K. A. (2023). Memory consolidation during sleep involves context reinstatement in humans. Cell Reports, 42(4), 112331. 10.1016/j.celrep.2023.112331

27. Schechtman, E., Witkowski, S., Lampe, A., Wilson, B. J., & Paller, K. A. (2020). Targeted memory reactivation during sleep boosts intentional forgetting of spatial locations. Scientific Reports, 10(1), 2327. 10.1038/s41598-020-59019-x Schreiner, T., Lehmann, M., & Rasch, B. (2015). Auditory feedback blocks memory benefits of cueing during sleep. *Nature Communications*, *6*(1), 8729.

28. Scotti, P. S., & Maxcey, A. M. (2022). Directed forgetting of pictures of everyday objects. Journal of Vision, 22(10), 8. 10.1167/jov.22.10.8

29. Simon, K. C. N. S., Gómez, R. L., & Nadel, L. (2018). Losing memories during sleep after targeted memory reactivation. Neurobiology of Learning and Memory, 151, 10–17. 10.1016/j.nlm.2018.03.003

30. Stickgold, R. (2005). Sleep-dependent memory consolidation. Nature, 437(7063), 1272– 1278. 10.1038/nature04286

31. Walker, M. P. (2009). The Role of Slow Wave Sleep in Memory Processing. Journal of Clinical Sleep Medicine, *5*(2 suppl). 10.5664/jcsm.5.2s.s20

32. Wernette, E. M. D., & Fenn, K. M. (2024). Factors influencing sleeplJdependent consolidation: Sleep strengths memory based on encoding depth but not repetition. Journal of Sleep Research, 33(5), e14091. 10.1111/jsr.14091

33. Whitmore, N. W., & Paller, K. A. (2023). Sleep disruption by memory cues selectively weakens reactivated memories. Learning & Memory, *30*(3), 63–69. 10.1101/lm.053615.122

34. Whitmore, N. W., Yamazaki, E. M., & Paller, K. A. (2024). Targeted memory reactivation with sleep disruption does not weaken week-old memories. Npj Science of Learning, 9(1), 64. 10.1038/s41539-024-00276-0

35. Woodward, A. E., & Bjork, R. A. (1971). Forgetting and remembering in free recall: Intentional and unintentional. Journal of Experimental Psychology, 89(1), 109–116. 10.1037/h0031188

36. Wylie, G. R., Foxe, J. J., & Taylor, T. L. (2008). Forgetting as an Active Process: An fMRI Investigation of Item-Method-Directed Forgetting. Cerebral Cortex, 18(3), 670–682. 10.1093/cercor/bhm101

37. Xia, T., Antony, J. W., Paller, K. A., & Hu, X. (2023). Targeted memory reactivation during sleep influences social bias as a function of slowlJoscillation phase and delta power. Psychophysiology, 60(5). 10.1111/psyp.14224

38. Yao, Z., Xia, T., Wei, J., Zhang, Z., Lin, X., Zhang, D., Qin, P., Ma, Y., & Hu, X. (2024). Reactivating cue approached positive personality traits during sleep promotes positive self-referential processing. iScience, 27(7), 110341. 10.1016/j.isci.2024.110341

