## Supplementary Materials for "Concurrent presentation of memory-related odors and sounds nullified sleep reactivation benefits"

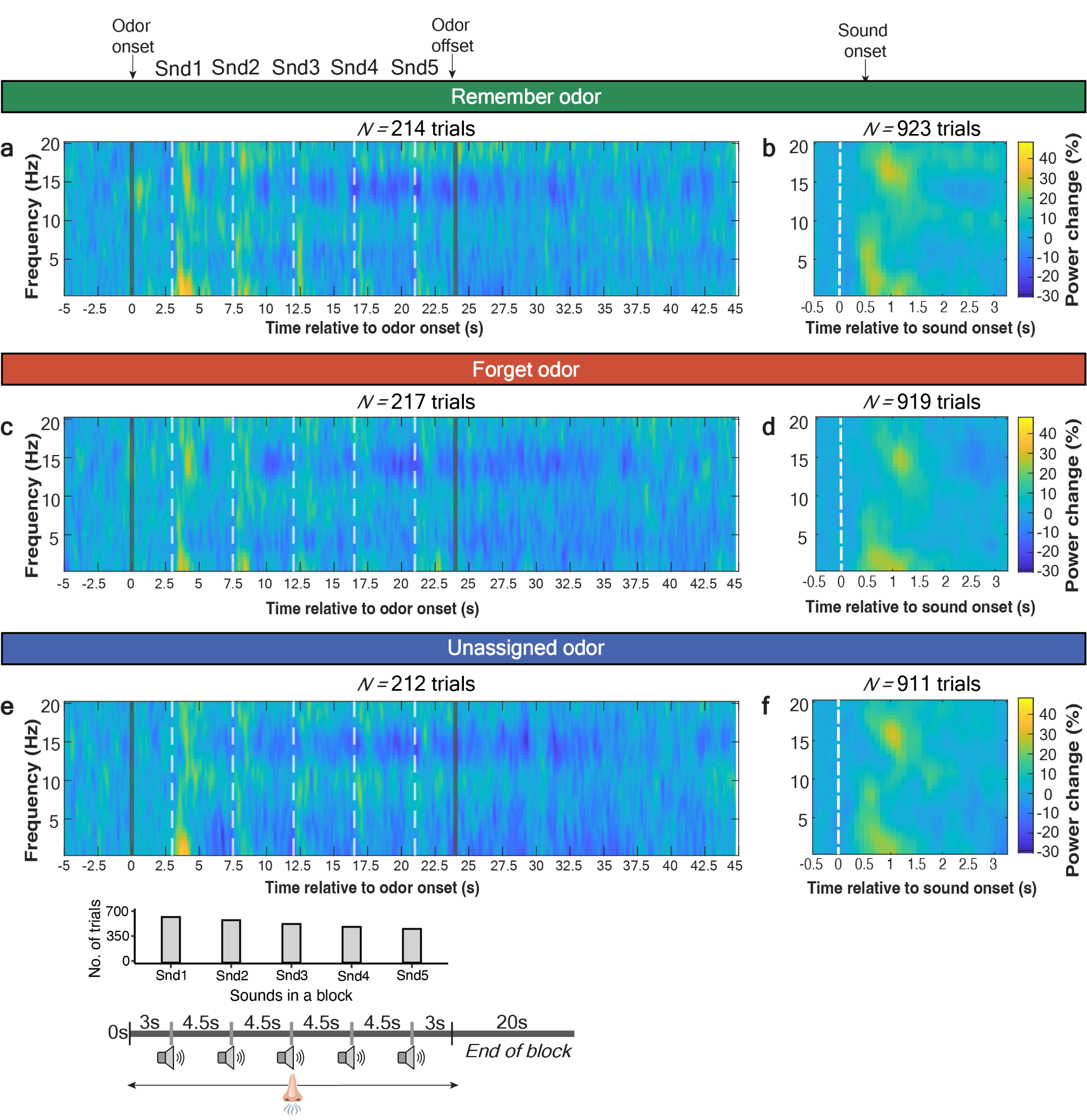

**Figure S1. Spectral response for all odors and sounds, regardless of sleep stage**

Baseline-corrected spectrograms displaying the percentage power change from baseline across the frequency bands (0–20 Hz) collapsed across participants. Gray lines indicate odor onset and offset. Snd1-Snd5 describes the onset of the five sounds presented with a 4.5-s onset-to-onset interval indicated by the white, dashed lines. The panels on the left column (**a,c,e**) describe activity in response to odor onset, and the panels on the right column (**b,d,f**) describe spectral activity in response to sound onset. The plot includes all trials without selecting based on NREM sleep. The trials include interrupted and uninterrupted cueing blocks.

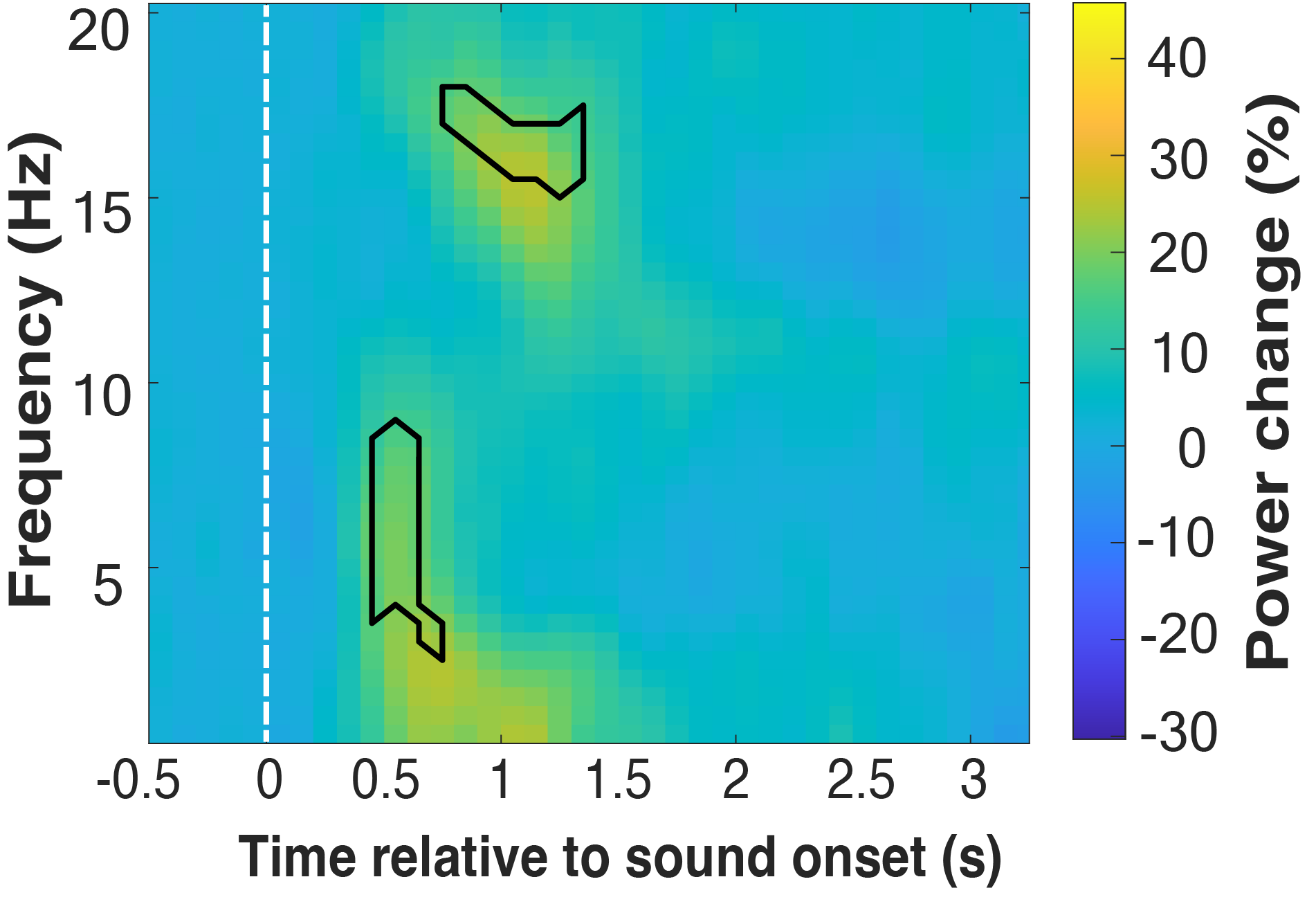

**Figure S2. Significant clusters of neural activity identified using a one-sample *t*-test**

Baseline-corrected spectrograms displaying the percentage power change from baseline across the frequency bands (0–20 Hz) collapsed across all participants and all sounds presented in NREM sleep. Sound onset is indicated by the white, dashed line. Significant clusters (α < .05) are indicated using the black lines and lie in the sigma and delta-theta range. Clusters were identified using a one-sample *t*-test across all sounds and participants, correcting for multiple comparisons.

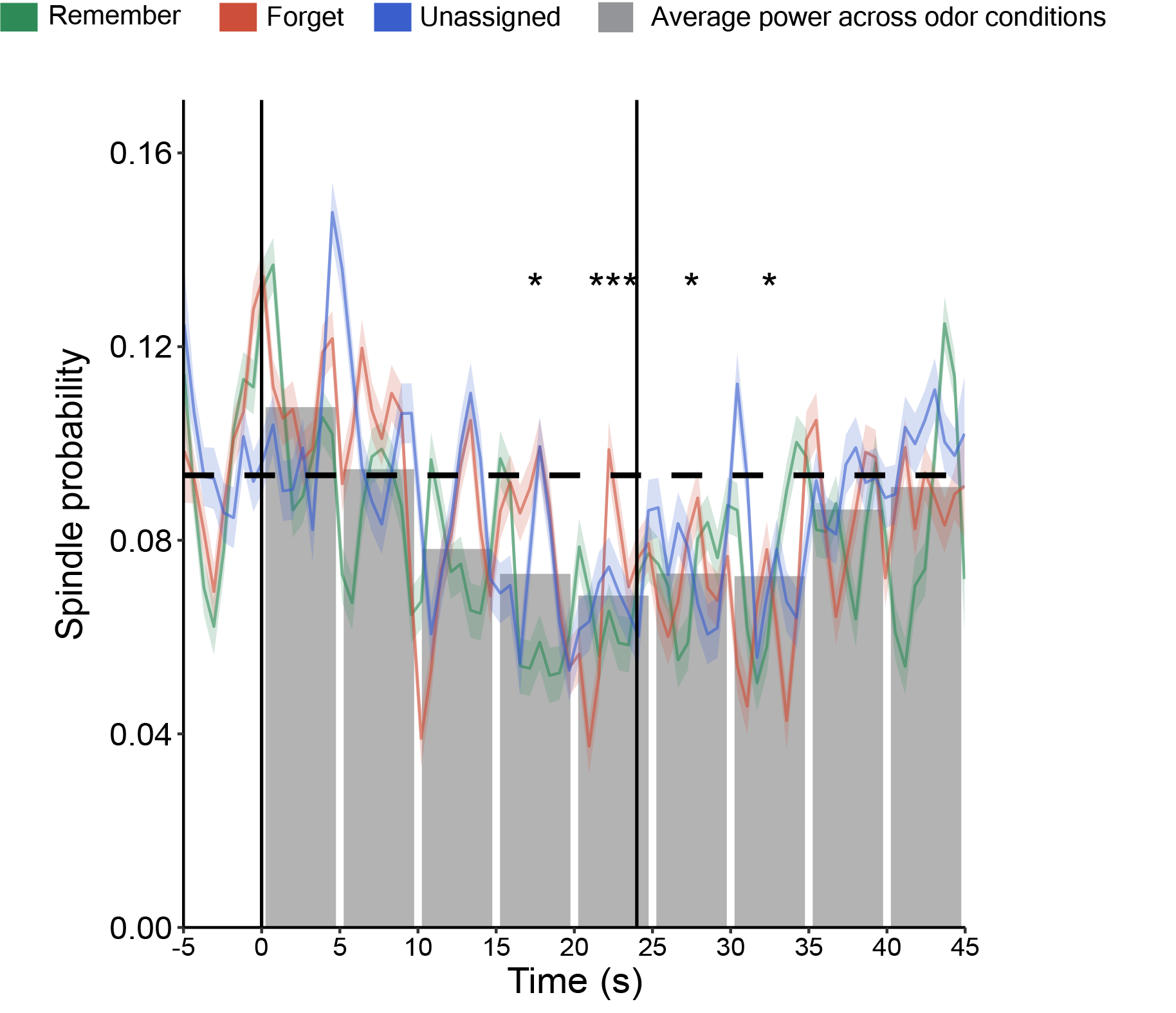

**Figure S3. Odor cues were followed by a prolonged decline in spindle probability.**

An algorithm was used to detect spindle occurrences at electrode Cz (e.g., Schechtman et al., 2021). Artifact-free EEG data were filtered between 11 and 16 Hz, and the root-mean-square (RMS) was calculated using a sliding window. Periods of time in which the RMS was higher than 1.5 times its standard deviation, lasting for 0.5–3 s, were considered as spindles. Next, for every odor trial, the number of spindle occurrences was computed throughout its duration. Collapsing this measure across trials yielded the temporal dynamics of spindle probability. We computed the spindle probability after odor onset across the three odor conditions, collapsed it across conditions, and averaged the data in 5-s epochs from 5 s before to 45 s after odor onset. Spindle occurrence probability of the 5-s epoch immediately before odor onset was considered as the baseline. A *t*-test was used to compare each 5-s epoch after odor onset to the baseline (corrected for multiple comparisons). Spindle occurrence probability declined significantly from 15 s after odor onset to 35 s after odor onset compared to baseline. The gray bars represent the averaged spindle probability in 5-s epochs. Black vertical solid lines indicate odor onset and offset. The dashed horizontal line indicates the baseline spindle probability. *p-*values were corrected for multiple comparisons. **p* < .05, ****p* < .001
